# A Bioinformatic Investigation into the Role of ITGB1 in Cancer Prognosis and Therapeutic Resistance

**DOI:** 10.1101/2025.04.06.647487

**Authors:** Xiaobin Mo

## Abstract

Integrin β 1 is a crucial transmembrane protein that regulates cellular adhesion, migration, and signal transduction, processes essential for cancer progression. This study investigates the role of ITGB1, the gene that encodes Integrin β1, in various cancers using bioinformatics tools. By analyzing gene expression data across different cancer types and normal tissues, the study identifies significant upregulation of ITGB1 in 12 cancer types. We find elevated expression of ITGB1 is associated with poor prognosis in multiple tumors, suggesting its potential as a biomarker for cancer progression and therapeutic resistance. Further analysis reveals ITGB1’s correlation with chemoresistance and immunoresistance genes, highlighting its involvement in cancer treatment evasion. The study also explores the expression and role of genes that are highly related to ITGB1 in tumor and patient prognosis, offering insights into potential molecular pathways and therapeutic targets. These findings underscore the clinical relevance of ITGB1 in cancer prognosis and therapy.

## Introduction

Studying the molecular mechanisms of genes in cancer requires an extensive exploration, as these mechanisms are influenced by a complex network of interactions [1], [2], [3], [4]. A comprehensive analysis of key genes involving major hallmarks of cancer [5], [6] across various tumor types can deepen our understanding and provide valuable insights into the development of targeted therapeutic strategies such as immune checkpoint blockades (ICBs) [7], [8], antibody-drug conjugates (ADC) [9], [10], and CAR-T therapy [11], [12]. Public databases such as GEPIA2, The Human Protein Atlas (THPA), and GeneCards enable bioinformatic analyses on large-scale patient database, facilitating the in-depth investigation of gene expression patterns, prognostic significance, and potential therapeutic implications in different cancer types [13], [14], [15]. These databases provide comprehensive resources and information for uncovering the multifaceted relationships between genes and cancer, offering a clearer perspective on their potential as biomarkers or therapeutic targets [16], [17].

The hallmarks of cancer define the biological capabilities acquired during tumor development, including self-sufficiency in growth signals, insensitivity to anti-growth signals, tissue invasion and metastasis, limitless replicative potential, sustained angiogenesis, evading apoptosis, etc. [5], [6]. Among these hallmarks, metastasis—the ability of cancer cells to spread from the primary tumor to distant organs—is particularly devastating, contributing to over 90% of cancer-related deaths [18]. Metastatic progression involves multiple complex processes, including epithelial-to-mesenchymal transition (EMT), extracellular matrix (ECM) remodeling, and the activation of signaling pathways that enhance cell motility and invasiveness [19], [20]. Given the critical role of metastasis in cancer lethality, focusing on key molecular players involved in this process is essential for understanding cancer progression and identifying potential therapeutic targets.

Integrin β1 (ITGB1) is a transmembrane receptor that regulates key signaling pathways involved in cell adhesion, migration, and survival [21]. Upon interaction with ECM components, ITGB1 activates focal adhesion kinase (FAK) and Src, modulating cytoskeletal dynamics and cell motility [22], [23], [24]. Additionally, ITGB1 signaling recruits PI3K and activates AKT, promoting cell proliferation and resistance to apoptosis [21]. In a tumor context, ITGB1 plays a key role in hepatocellular carcinoma (HCC) by activating the FAK/AKT signaling pathway, promoting cell proliferation and invasion [25]. Another study show that rAj-Tspin, a novel peptide from *A. japonicus*, suppresses ITGB1 expression and inhibits the FAK/AKT pathway in Huh7 hepatocellular carcinoma cells. It further reducing expression of EMT markers and promoting apoptosis, which hinders hepatocellular tumor progression [26]. In gastric cancer, high ITGB1 expression is correlated with poor prognosis [27]. In breast cancer, linc-ITGB1 promotes cancer progression by inducing cell cycle arrest and interrupting the EMT process [28]. In ovarian cancer, cancer-associated fibroblasts expressing DDR2 promote tumor invasion and metastasis through Periostin-ITGB1 [29].

Despite the growing recognition of ITGB1’s role in cancer, its bioinformatic analysis in cancer prognosis remains less well-established. This study aims to analyze ITGB1’s expression patterns and prognostic significance using bioinformatics tools. We first investigated the expression of ITGB1 in multiple tumor types at varied stages (Figures 1, 2). Then, we conducted survival analysis of ITGB1 expression across 33 cancer types and identified specific cancers whose survival rates showed significant correlation with ITGB1 expression changes. Next, we performed a correlation analysis between ITGB1 and major chemoresistance and immunoresistance genes (Figures 4, 5). Finally, we identified ten genes associated with ITGB1 and examined their significance in cancer prognosis (Figures 6, 7).

## Results

### ITGB1 Structure, Localization, and Expression Across 20 Cancer Types

Using AlphaFold, we generated the predicted visualization of the three-dimensional structure of ITGB1 (Figure 1a). To assess the localization of ITGB1 in cells, we obtained fluorescence images of ITGB1 from THPA for three cell lines. Immunofluorescence images verified the expression and subcellular localization of ITGB1 in multiple tumor types including in the epidermoid carcinoma A-431 cell line, the osteosarcoma U2OS cell line, and the malignant glioblastoma U-251MG cell lines. To analyze the expression level of ITGB1 in multiple tumor sites, we first obtained the bodymap showing the median expression levels of ITGB1 in tumor and normal tissues from GEPIA2 (Figure 1c). The bodymap showed overexpression of ITGB1 (indicated by darker color) in some tumor samples (red) compared to normal tissues (green) across various cancer types. Additionally, we analyzed data from THPA showing the protein-level across 20 cancer types (Figure 1d). On average, more than 90% of patients show high or medium expression of ITGB1 across these 20 cancer types, indicating that ITGB1 expression was a tumor signature in multiple tumor types.

**Figure 1.**
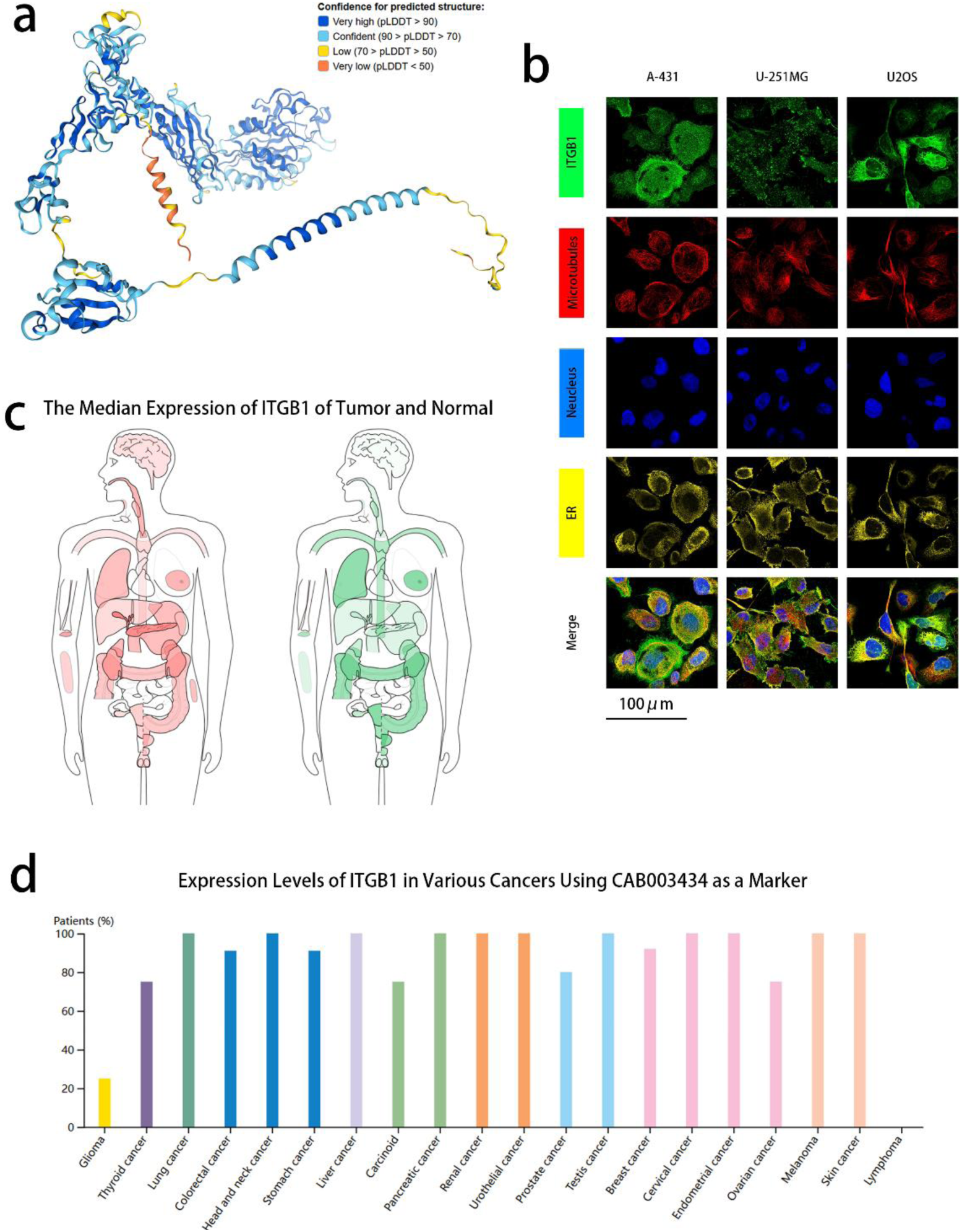
(a) Protein Structure Visualization from THPA. The model predicted the complete structure of the ITGB1 protein. Different colors in the structural model represent the prediction confidence. (b) Subcellular localization images from THPA. These three cells are derived from the A-431, the U-251MG and the U2OS cell line respectively, and their antibody catalog number is CAB003434. Through fluorescence microscopy, the localization of the target protein ITGB1 in the endoplasmic reticulum and plasma membrane can be observed. The image also shows the structures of the nucleus and microtubules. The green color represents the target protein ITGB1, the red color represents the microtubules, the blue color represents the nucleus, and the yellow color represents the ER. (c) The median expression of ITGB1 of tumor and normal samples in bodymap from GEPIA2. The red on the left represents tumor, and the green on the right represents normal. (d) The protein expression level of ITGB1 in 20 cancer types from THPA.

### ITGB1 expression is upregulated in 12 cancer types

To quantitatively investigate the expression level of ITGB1, we compared its expression (log2(TPM+1) values) in tumor samples from 31 TCGA cancer types against a combined normal tissue dataset (integrating TCGA normal and GTEx samples). Analysis of gene expression data from GEPIA2 revealed that ITGB1 was significantly upregulated in CHOL, DLBC, ESCA, GBM, HNSC, LGG, PAAD, PCPG, SKCM, STAD, TGCT, and THYM. The log10(T/N) heatmap illustrates this widespread overexpression in these cancer types (Figure 2a), where “T” represents tumor data from TCGA, and “N” denotes the matched normal tissue data. The boxplot analysis further provide details of individual samples and the differences between tumor and normal tissues in the upregulated tumor types (Figure 2a). The increased expression of ITGB1 in these cancers suggests a potential role in tumorigenesis, possibly through promoting cell adhesion, migration, and survival, which are critical processes in cancer progression. In addition to the overexpression analysis, we also examined ITGB1 expression across different cancer stages to explore its potential as a biomarker for disease progression. In four cancer types (ACC, BLCA, LIHC, SKCM), we observed a significant increase in ITGB1 expression as the cancer stage advanced (Figure 2b). This suggests that ITGB1 may not only be involved in the initiation of cancer but also in the progression of the disease, highlighting its potential as a therapeutic target in later stages of cancer.

**Figure 2.**
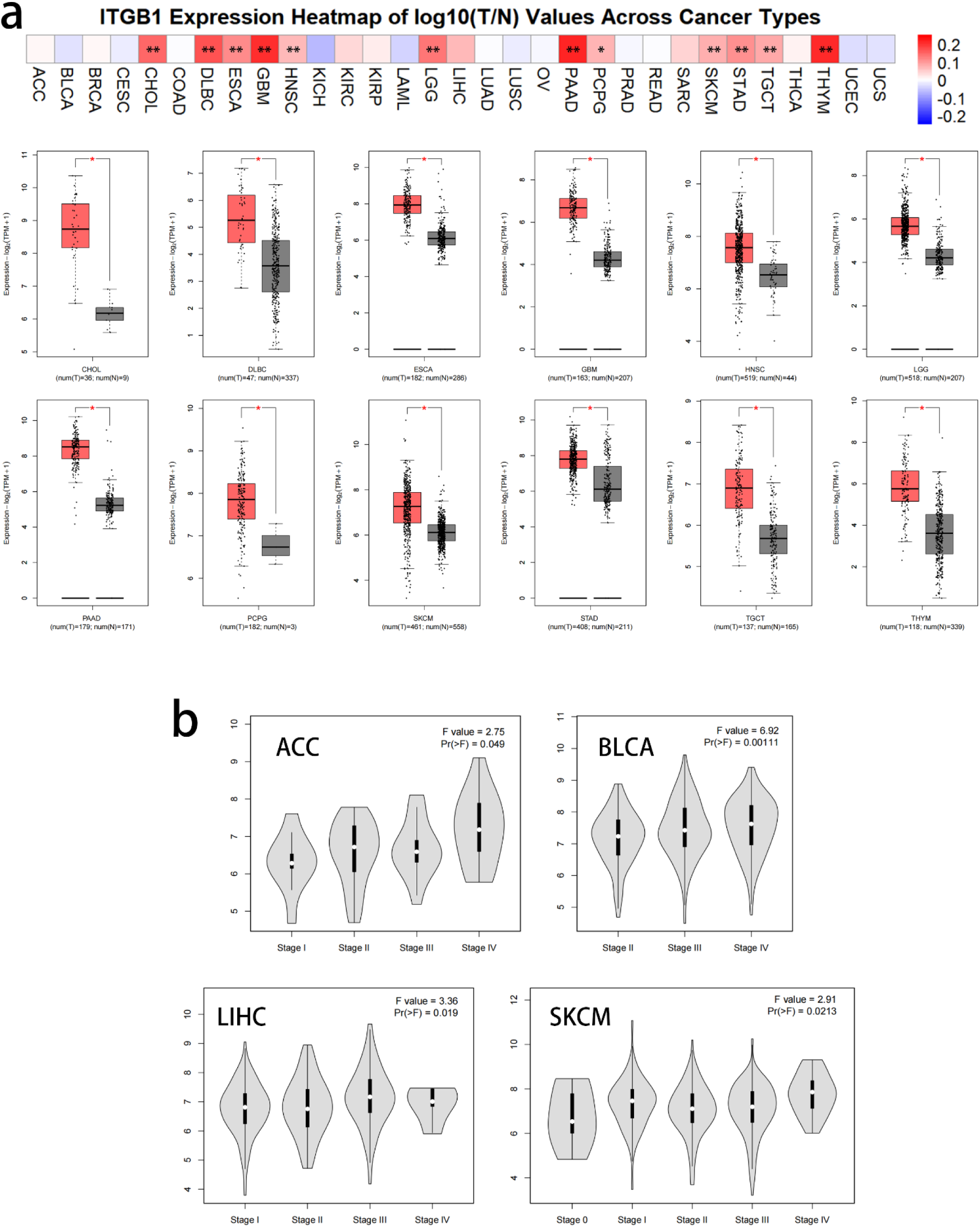
(a) Heatmap showing the log2 fold-change of ITGB1 expression (Tumor/Normal) across 31 cancer types, and box plots displaying the differential expression (log2(TPM+1) values) of ITGB1 between normal and tumor tissues in 12 cancer types where ITGB1 is significantly upregulated. The data are from GEPIA2. (b) Expression levels of ITGB1 (log2(TPM+1)) at different pathological stages of 4 cancer types where ITGB1 showed significant expression differences across different stages. *P < 0.05.

### Elevated levels of ITGB1 indicate poorer survival rates in various cancers

To investigate the potential relationship between ITGB1 expression and patient prognosis across various cancer types, we conducted survival analysis for 33 cancer types. We plotted the survival graphs showing the survival outcomes of patients separated by high and low ITGB1 expression (Figure 3), in which we observed significant differences in Overall Survival (OS) and Disease Free Survival (DFS) outcomes associated with ITGB1 expression in multiple tumor types. In terms of OS, high ITGB1 expression correlated with shorter overall survival rates in cancers such as ACC, CESC, KIRP, LGG, LIHC, LUAD, LUSC, MESO, PAAD, and STAD (Figure 3a). Similarly, for DFS, increased ITGB1 expression was linked to poorer disease-free survival in cancers including ACC, CESC, COAD, LGG, LUAD, LUSC, MESO, PAAD, and SARC (Figure 3b). The analysis revealed that elevated ITGB1 expression was significantly associated with poorer survival outcomes across several cancer types. The observed correlation between high ITGB1 expression and worse survival outcomes could reflect its involvement in processes such as tumor invasion, metastasis, and resistance to treatment. Elevated ITGB1 levels may promote cell adhesion and migration, which are critical for cancer cells to invade surrounding tissues and establish secondary tumors. These findings suggest that ITGB1 may serve as a prognostic marker for poor outcomes, indicating its potential as a target for improving patient outcomes.

**Figure 3.**
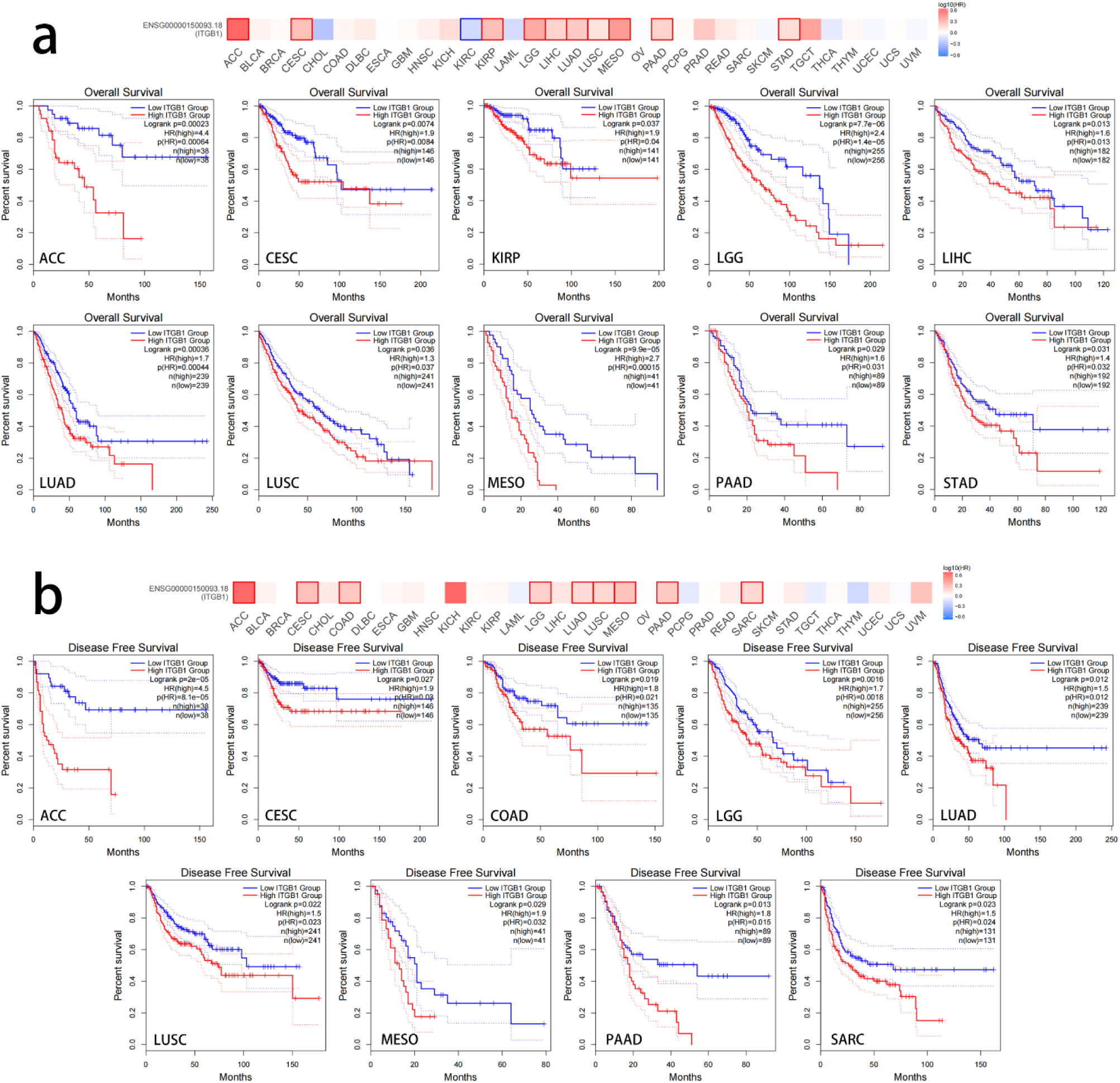
Correlation between ITGB1 expression and patient prognosis across 33 cancer types based on GEPIA2 data. The heat maps and survival curves showing significant differences in overall survival (a) and disease-free survival (b) for different cancer types. Highlighted outlines in heatmaps indicate signicant differences. The red survival curves represent high ITGB1 groups, and the blue survival curves represent low ITGB1 groups. Elevated ITGB1 expression was associated with worse OS in 10 cancer types and with poorer DFS in 9 cancer types.

### Correlation Between ITGB1 Expression and Chemoresistance Genes Across 33 Cancer Types

To investigate the potential association between ITGB1 expression and chemoresistance-related genes, we focused on 10 key genes that are implicated in drug resistance mechanisms. We analyzed the correlation between ITGB1 expression and 10 chemoresistance-related genes (ABCB1, ABCC1, ABCG2, BCL2, BAX, CASP3, ERCC1, MGMT, TOP2A, and RRM1) [30], [31], [32], [33], [34], [35], [36] and calculated the Pearson Correlation Coefficient (PCC). Then, we plotted a heat map to visualize these correlations (Figure 4a). Overall, we detected a high percentage of significant positive correlation between ITGB1 and the chemoresistance genes across the tumor types. Higher ITGB1 expression was positively correlated with the expression of chemoresistance-related genes such as ABC transporters (ABCB1, ABCC1, and ABCG2), anti-apoptotic genes (BCL2, CASP3), and DNA repair genes (TOP2A, RRM1) (Figure 4a). Specifically, the numbers of genes positively correlated with ITGB1 among the 10 chemoresistance-related genes are highest in LGG, KIRP, OV and UVM across 33 cancer types. LGG, KIRP, OV, and UVM each exhibit a high number of correlation between ITGB1 and chemoresistance-related genes, with 9, 9, 8 and 7 out of 10 chemoresistance-realated genes respectively (Figure 4b). This suggests a stronger correlation between ITGB1 expression and chemoresistance in multiple cancer types. In these 10 chemoresistance-related genes, CASP3, ABCC1, and RRM1 exhibit the highest positive correlation with ITGB1 across cancer types, correlating in 29, 27, and 26 out of 33 cancers, respectively (Figure 4c). These findings suggest that ITGB1 may play a role in promoting chemoresistance in tumors.

**Figure 4.**
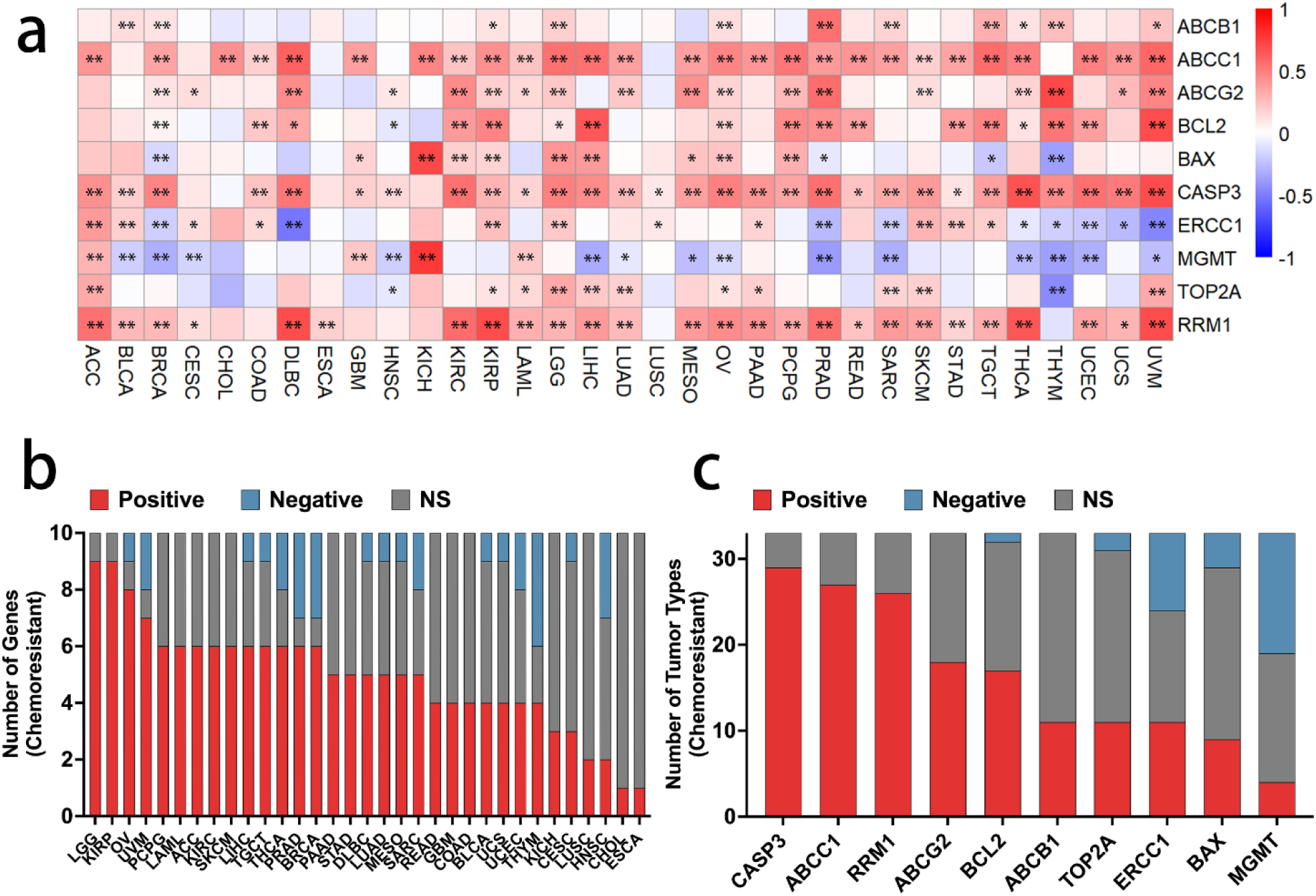
(a) Heatmap of the correlation (PCC) between ITGB1 and chemoresistance-related genes in 33 cancer types. (b) Heatmap of the correlation (R-value) between ITGB1 and immunoresistance-related genes. (b) Histogram showing the number of genes that are positively correlated, negatively correlated, or non-significant with ITGB1 among the 10 chemoresistance-related genes across 33 cancer types. (c) Histogram showing the number of cancer types in which the 10 chemoresistance-related genes are positively correlated, negatively correlated, or non-significant with ITGB1.

### Correlation Between ITGB1 Expression and Immunoresistance Genes Across 33 Cancer Types

We further investigated the potential relationship between ITGB1 expression and immunoresistance-related genes to assess its role in immune evasion in cancer. We analyzed the correlation between ITGB1 expression and 10 major immunoresistance-related genes (PD-1, PD-L1, CTLA-4, LAG3, TIM-3, TIGIT, TGFB1, IDO1, CD47, and ARG1) [37], [38], [39], [40], [41], [42] and calculated the PCC. A heatmap was generated to visualize these correlations (Figure 5a). We also detected a high percentage of significant positive correlation between ITGB1 and the immunoresistance genes across the tumor types (Figure 5a). The numbers of genes positively correlated with ITGB1 among the 10 immunoresistance-related genes are highest in LGG, OV, PCPG, BLCA and PRAD across 33 cancer types. LGG, OV, PCPG, BLCA and PRAD each exhibit a high number of correlation between ITGB1 and immunoresistance-related genes, with 9, 8, 8, 8 and 8 out of 10 immunoresistance-related genes respectively (Figure 5b). This suggests a stronger correlation between ITGB1 expression and immunoresistance in these tumor types. In these 10 immunoresistance-related genes, TGFB1, CD47, and TIM-3 exhibit the highest positive correlation with ITGB1 across cancer types, correlating in 27, 21 and 19 out of 33 cancers, respectively (Figure 5c). Our analysis revealed significant positive correlations between ITGB1 expression and several immunoresistance-related genes across different cancer types. Specifically, ITGB1 expression showed significant positive correlations with immune evasion markers including PD-L1, CTLA-4, TIM-3, TIGIT, TGFB1, and CD47. These findings suggest that ITGB1 may play a role in driving immunoresistance in tumors.

**Figure 5.**
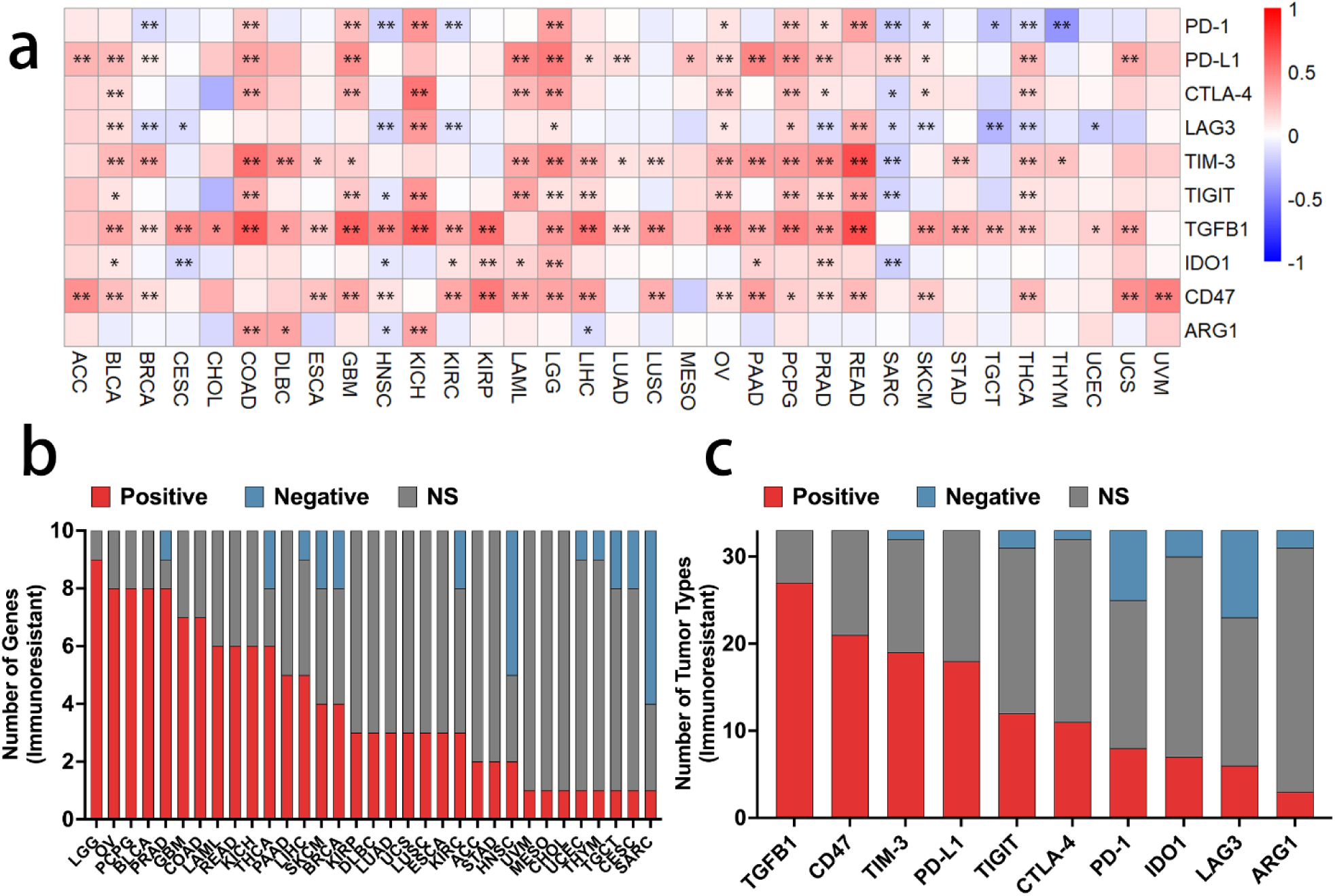
(a) Heatmap of the correlation (PCC) between ITGB1 and immunoresistance-related genes in 33 cancer types. (b) Histogram showing the number of genes that are positively correlated, negatively correlated, or non-significant with ITGB1 among the 10 immunoresistance-related genes across 33 cancers types. (c) Histogram showing the number of cancer types in which the 10 immunoresistance-related genes are positively correlated, negatively correlated, or non-significant with ITGB1.

### Identification of Genes that are Highly Correlated to ITGB1 in Multiple Cancer Types

To explore the role of ITGB1-related genes across different cancer types and their impact in cancer prognosis, we conducted STRING analysis to identify the top 25 genes and top 5 genes (LAMC1, ITGA6, FN1, ITGA5, ITGB1BP1) that are strongly correlated with ITGB1 (Figure 6a). Additionally, using the Similar Genes Detection feature of GEPIA2 as a separate independent analysis, we further identified another five genes with the highest PCC with ITGB1 (ITGB1P1, SEC23A, COL4A1, MYH9, COL4A2) (Figure 6b), making a gene set containing a total of 10 highly related genes. We plotted the log10(T/N) expression heatmap of the 10 genes across different cancer types (Figure 6c). The heatmap reflects that these genes are also mostly upregulated in multiple tumor types (Figure 6a), which suggests genes that are highly related to ITGB1 are also associated with tumorigenesis.

**Figure 6.**
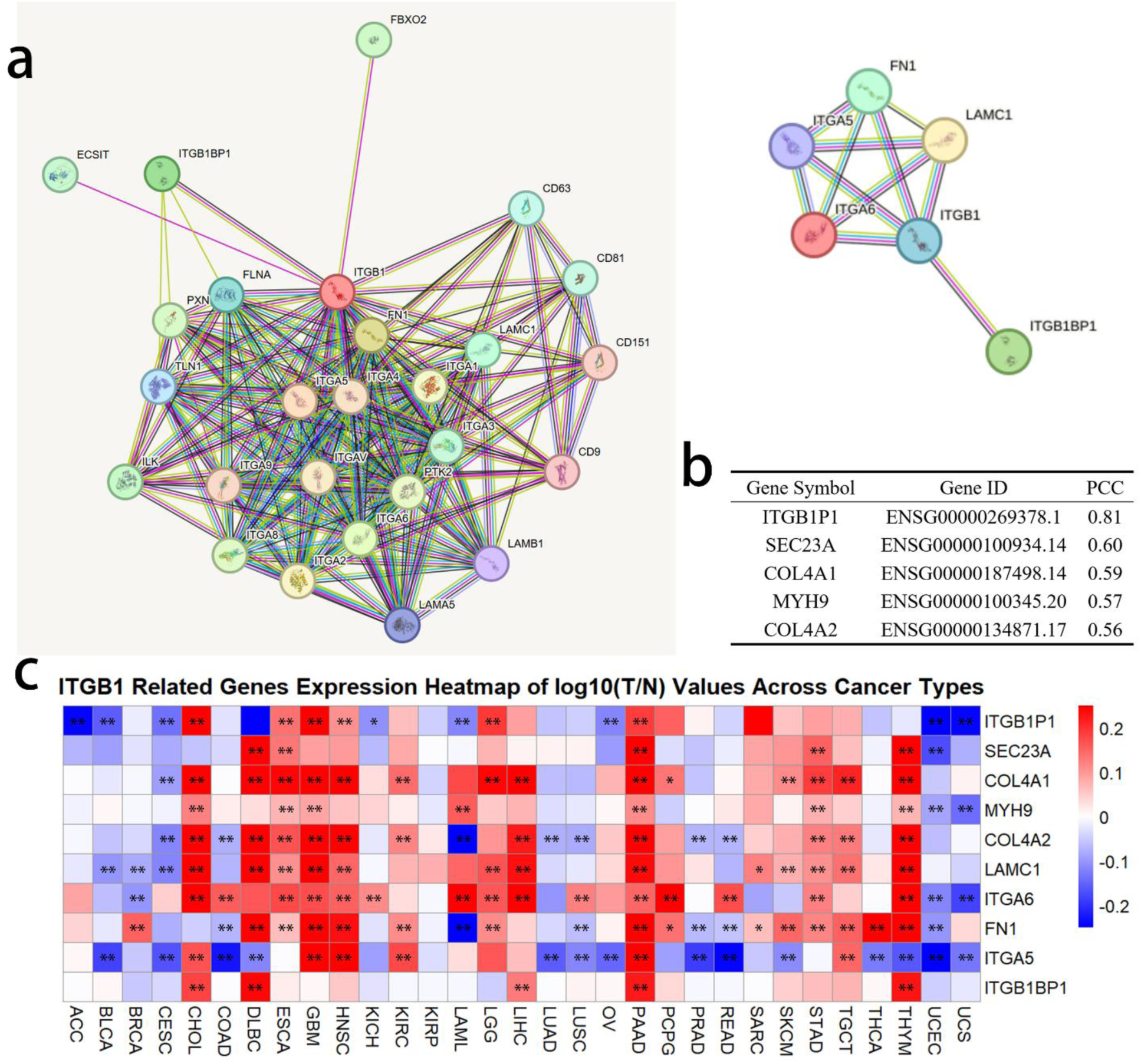
(a) The top 25 genes and top 5 genes most related to ITGB1 in the String analysis on GeneCards. (b) The five genes with the highest PCC with ITGB1 across all TCGA Tumor data using the Similar Genes Detection feature on GEPIA2. (c) The expression heatmap of log10(T/N) values for these 10 genes across cancer types. The data are from GEPIA2.

### Overall Survival Analysis of Genes Related to ITGB1

To assess the role of ITGB1-related genes in patient outcomes, we plotted the OS heatmap for these 10 genes across different cancer types (Figure 7a). The analysis shows that for those that are significantly affected, the patients survival outcomes are mostly getting worse when there is overexpression of these ITGB1-related genes. In 33 cancer types, the highest number of genes related to ITGB1, whose high expression is significantly associated with worse OS, are found in LGG, MESO, and UVM, with 10, 8, and 6 out of 10 genes, respectively (Figure 7b). Among these 10 ITGB1-related genes, the high expression of ITGA5, COL4A1, and FN1 is significantly linked to worse OS in the most cancer types, with 11, 9, and 8 out of 33, respectively (Figure 7c).

**Figure 7.**
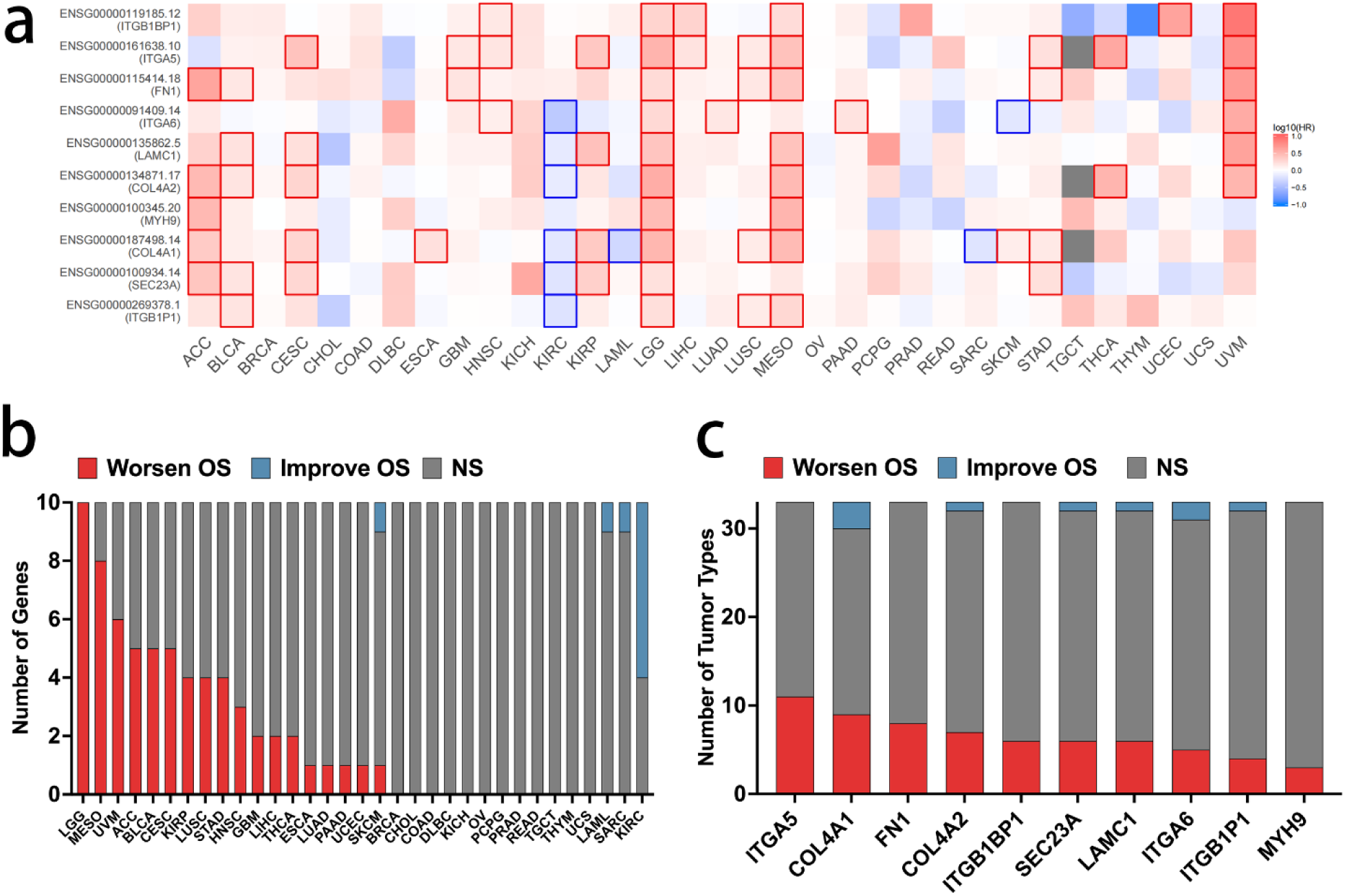
(a) The overall survival heatmap of these 10 genes across cancer types. Highlighted outlines indicate significant differences. The data are from GEPIA2. (b) A histogram representing the number of genes among these 10 ITGB1-related genes whose high expression is associated with worsening OS, improving OS, or showing no significance across various cancer types. (c) A histogram representing the number of cancer types for each of these 10 ITGB1-related genes where high expression is associated with worsening OS, improving OS, or showing no significance.

### Disease-Free Survival Analysis of Genes Related to ITGB1

We also plotted the DFS heatmap for these 10 genes across different cancer types (Figure 8a), with data from GEPIA2. Like OS analysis, DFS analysis also shows that the expression levels of several of these genes significantly influence patients’ DFS survival, with higher expression mostly correlated with poorer survival outcomes in many cancers. The highest number of genes related to ITGB1, whose high expression is significantly associated with worse DFS, are found in ACC, LGG, and UVM, with 8, 7, and 6 out of 10 genes, respectively (Figure 8b). Among these 10 ITGB1-related genes, the high expression of ITGA5, ITGB1BP1, and FN1 is significantly linked to worse DFS in the most cancer types, with 8, 8, and 6 out of 33, respectively (Figure 8c).

**Figure 8.**
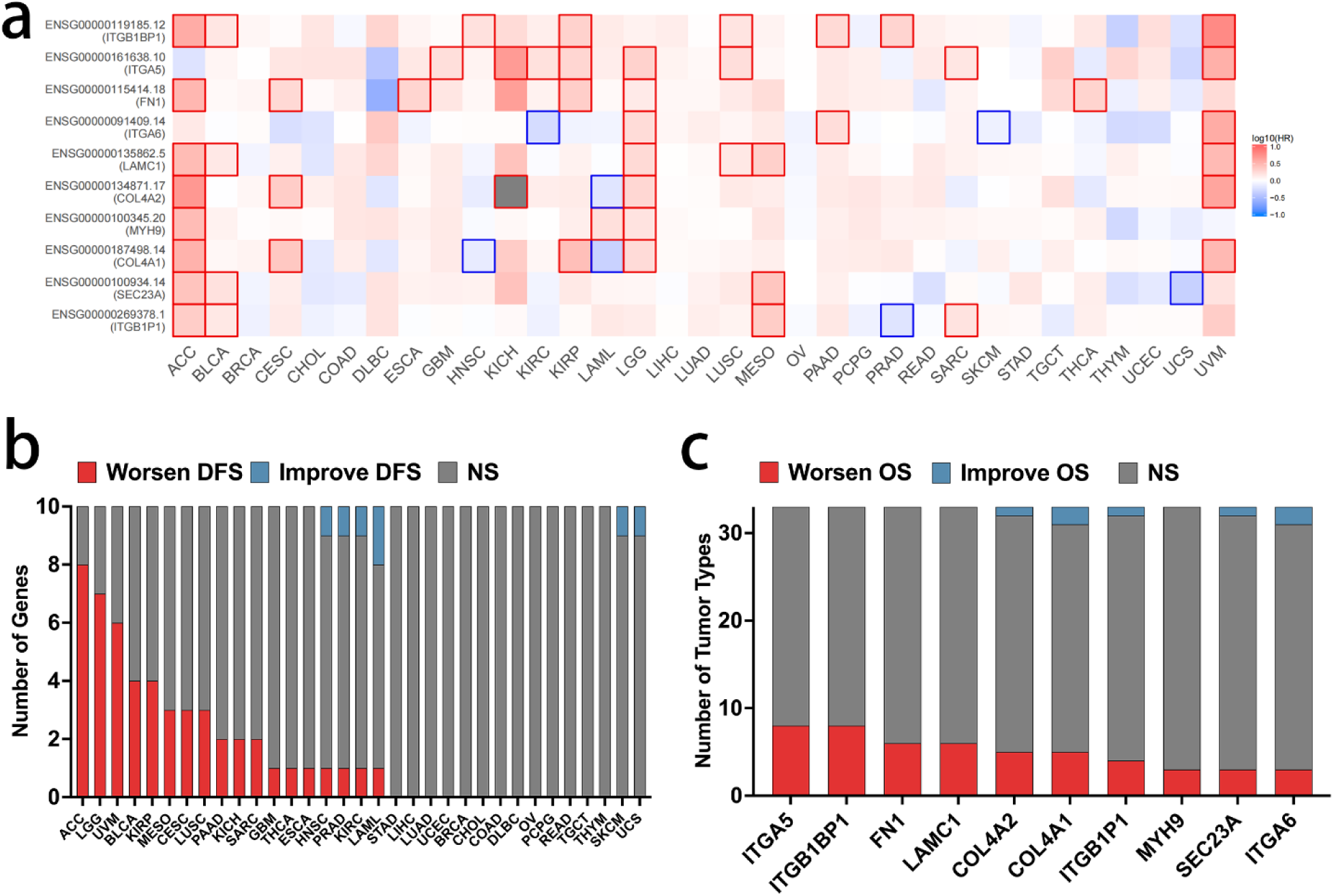
(a) The disease-free survival heatmap of these 10 genes across cancer types. Highlighted outlines indicate signicant differences. The data from GEPIA2. (b) A histogram representing the number of genes among these 10 ITGB1-related genes whose high expression is associated with worsening DFS, improving DFS, or showing no significance across various cancer types. (c) A histogram representing the number of cancer types for each of these 10 ITGB1-related genes where high expression is associated with worsening DFS, improving DFS, or showing no significance.

## Discussion

### ITGB1 as a Clinical Marker: Implications for Cancer Prognosis

ITGB1 is a key transmembrane protein that plays a critical role in cellular adhesion, migration, and signal transduction, all of which are essential processes in cancer progression [43], [44], [45]. Our findings demonstrate that the upregulation of ITGB1 directly correlates with poor patient prognosis, making ITGB1 a promising clinical biomarker for cancer progression (Figures 2, 3). This correlation is particularly evident in cancers such as ACC, CESC, KIRP, LGG, and LUAD, where elevated ITGB1 expression is linked to shorter overall survival and disease-free survival times (Figure 3). Elevated ITGB1 levels likely contribute to tumor aggressiveness by promoting cell adhesion and migration, key processes for metastasis. There are studies show that ITGB1 interacts with ECM components such as fibronectin and collagen at the cell membrane, forming focal adhesion complexes that anchor cells within their microenvironment [22], [23]. Upon activation, ITGB1 stimulates FAK, which subsequently activates Src and regulates cell motility, with paxillin orchestrating these signals at focal adhesions [24]. This dynamic interaction plays a key role in cell migration by enabling integrin-mediated adhesion turnover, facilitating the detachment and reattachment of cancer cells during movement [46]. Furthermore, ITGB1 promotes cell survival, proliferation, and resistance to anoikis by inducing FAK autophosphorylation, recruiting PI3K, and activating AKT [21]. Moreover, high ITGB1-DT expression activates the MAPK/ERK signaling pathway, linked to cell proliferation, migration, and invasion. Interfering with ITGB1-DT expression reduces the phosphorylation of p38 MAPK and ERK, thus limiting cancer cell proliferation and migration [47].

Furthermore, ITGB1’s association with cancer progression suggests its potential utility as a therapeutic target, particularly in late-stage cancers where its expression is significantly increased. The combination of high ITGB1 expression with poor survival outcomes underscores its role as an important prognostic marker in various cancers (Figure 3). By influencing processes like EMT and cell survival, ITGB1 enhances tumor invasion and metastasis [48], [49], which are often the primary causes of poor prognosis. As a result, ITGB1’s overexpression serves as a signal of aggressive tumor behavior, indicating that targeting ITGB1 could help mitigate these effects, providing new avenues for cancer treatment strategies. This role as a clinical marker highlights the need for further exploration of ITGB1 as a prognostic tool, potentially allowing for the development of novel ITGB1-targeted therapies, such as ADC and CAR-T therapies that targets the integrin β1.

### Association with Chemoresistance and Immunoresistance: A Key Player in Therapeutic Resistance

Our analysis also reveals that ITGB1 is intricately associated with chemoresistance and immunoresistance genes, suggesting its involvement in therapeutic resistance mechanisms. Chemoresistance, a major challenge in cancer treatment, is often mediated by the upregulation of certain resistance genes that enable tumor cells to evade the effects of chemotherapy [50]. In our study, ITGB1 expression was positively correlated with several key chemoresistance-related genes (Figure 4), including ABC transporters (ABCB1, ABCC1, ABCG2) [51], anti-apoptotic genes (BCL2, CASP3) [52], [53], and DNA repair genes (TOP2A, RRM1) [54], [55]. These genes are known to confer resistance to chemotherapy drugs by either pumping them out of cancer cells, inhibiting apoptosis, or enhancing DNA repair mechanisms that allow cancer cells to survive after treatment. Our analysis identified the highest number of genes positively correlated with ITGB1 among the 10 chemoresistance-related genes are found in LGG, KIRP, OV, and UVM (Figure 4b). This suggests a stronger link between ITGB1 expression and chemoresistance in these tumors. Across these 33 cancer types, CASP3, ABCC1, and RRM1 show the strongest correlation with ITGB1 (Figure 4c), suggesting they may play a key role in ITGB1-mediated chemoresistance.

The link between ITGB1 and these chemoresistance genes may suggests that ITGB1 may act as a facilitator of therapeutic resistance, potentially by modulating cellular adhesion and survival pathways that enable cancer cells to resist chemotherapy. For instance, ITGB1’s role in the FAK/AKT and MAPK/ERK pathways may enhance cell survival and proliferation even in the presence of chemotherapeutic agents [56], [57], [58]. This reinforces the clinical significance of ITGB1 as a potential target to overcome chemoresistance, offering new strategies for improving the efficacy of existing therapies.

Similarly, ITGB1 is also implicated in immunoresistance, a phenomenon where cancer cells evade detection and destruction by the immune system. We found that ITGB1 expression was positively correlated with several immune checkpoint genes (Figure 5), such as PD-L1, CTLA-4, TIM-3, and TIGIT. These immune checkpoints play critical roles in suppressing anti-tumor immunity, allowing cancer cells to avoid immune-mediated destruction [59], [60], [61]. For example, through interaction with the PD-1 receptor on T cells, PD-L1 inactivates these immune effectors [62]. Our analysis identified the highest number of genes positively correlated with ITGB1 among the 10 immunoresistance-related genes are found in LGG, OV, PCPG, BLCA, and PRAD (Figure 5b). This suggests a stronger link between ITGB1 expression and immunoresistance in these tumors. TGFB1, CD47, and TIM-3 show the strongest correlation with ITGB1 (Figure 5c), indicating they may play a central role in ITGB1-mediated immunoresistance. With future verification through gene knockdown, ITGB1 may enhance the ability of cancer cells to escape immune surveillance by promoting the expression of these immune evasion markers, contributing to their resistance to immunotherapy. The link between ITGB1 and immune resistance genes suggests that targeting ITGB1 may not only improve chemotherapy response but also enhance the effectiveness of immunotherapies, particularly immune checkpoint inhibitors.

### Correlation Analysis: High-Correlated Genes with ITGB1

The correlation analysis conducted in this study shows the important role of ITGB1 in cancer progression, highlighting its interaction with key genes and pathways. These genes are integral to critical cellular processes such as cell adhesion, migration, survival, and immune modulation, all of which contribute to tumorigenesis and metastatic spread (Figure 6).

Among the top genes correlated with ITGB1, several are known to play key roles in cell motility, invasion, and survival—critical aspects of cancer biology. For instance, genes like LAMC1, ITGA6, FN1, and ITGA5 are involved in cellular adhesion and migration [63], [64], [65], [66], while ITGB1BP1 further mediates interactions between integrins and the ECM, contributing to the metastatic potential of tumor cells [67], [68]. These interactions suggest that ITGB1 not only facilitates cancer cell adhesion but also enhances the ability of tumor cells to migrate and invade surrounding tissues, thereby promoting metastasis.

Furthermore, the association between ITGB1 and genes involved in immune response pathways, such as SEC23A and COL4A1, suggests its involvement in immune evasion. In particular, COL4A1 and COL4A2 are known to regulate ECM remodeling, a process that can alter immune cell infiltration and hinder effective immune surveillance [69]. By modulating the tumor microenvironment in this manner, ITGB1 may help cancer cells evade immune detection, thereby contributing to both tumor progression and resistance to immune-based therapies.

In addition to immune evasion, our findings highlight ITGB1’s potential role in chemoresistance. Several studies suggest a relationship between FN1, ITGA6, and chemoresistance in various cancers [70], [71]. Our study suggests that ITGB1, through its interactions with genes such as FN1 and ITGA6, could modulate the tumor’s response to chemotherapy. These findings reinforce the idea that ITGB1 is not only a key player in cancer progression but also in therapeutic resistance, particularly in the context of chemotherapy and immunotherapy.

The highest number of ITGB1-related genes associated with worse OS are found in LGG, MESO, and UVM (Figure 7b). This suggests a stronger link between ITGB1 expression and poor prognosis in these tumor types. ITGA5, COL4A1, and FN1 show the strongest correlation with ITGB1 in OS (Figure 7c), indicating their potential role in ITGB1-mediated metastasis, immunoresistance and chemoresistance.

The highest number of ITGB1-related genes associated with worse DFS are found in ACC, LGG, and UVM. This suggests a stronger link between ITGB1 expression and poor prognosis in these tumor types (Figure 8b). ITGA5, ITGB1BP1, and FN1 show the strongest correlation with ITGB1 in DFS (Figure 8c), suggesting their involvement in ITGB1-mediated recurrence and progression.

## Methods

### Protein Structure Visualizations and Subcellular localization image

The protein structure of ITGB1 was obtained from THPA. Specifically, the structure was visualized using the AlphaFold v2.3.2 model, which provides some predictions of protein structures based on deep learning techniques. Three subcellular localization fluorescence images of ITGB1 were obtained from THPA. Using fluorescence microscopy, the localization of the target protein ITGB1 was observed in the plasma membrane.

### Comprehensive Analysis of ITGB1 Expression Across Organs, Cancer Types, and Stages

The expression levels of ITGB1 in various organs from both cancer and normal tissues were analyzed using GEPIA2. A protein expression map across different cancers was obtained from THPA. The data were used to compare the relative expression of ITGB1 between different organs, providing a comprehensive understanding of its expression patterns in healthy versus tumor tissues.

Gene expression data for ITGB1 across 31 cancer types and normal tissues were obtained from GEPIA2. A log10(T/N) heatmap was generated using the pheatmap package in R to illustrate the expression differences between tumor and normal tissues (Figure 2a), where “T” denotes tumor data from TCGA, and “N” is a combination of TCGA normal samples and GTEx samples. Additionally, a boxplot was created for cancers showing statistically significant differences in ITGB1 expression from GEPIA2, where the sample sizes for T and E in each cancer type are annotated below the corresponding boxplot. (Figure 2a).

Using GEPIA2, stage plots were generated to assess how ITGB1 expression varies across different stages of cancer development. The analysis specifically focused on cancer stages with significant expression differences (Figure 2b).

### Survival Analysis

The impact of ITGB1 expression on patient survival was assessed using GEPIA2. A heatmap for both OS and DFS was created from GEPIA2, and the corresponding Kaplan-Meier survival curves were plotted for cancers where significant differences in survival were observed based on ITGB1 expression levels (Figure 3). The group cutoff was set to the median, and the sample sizes for the high- and low-expression groups are annotated on the survival curves.

### Chemoresistance and Immunoresistance Correlation

Gene expression data from GEPIA2 were used to analyze the correlation between ITGB1 and known chemoresistance and immunoresistance genes. Two heatmaps were constructed using the pheatmap package in R to illustrate the relationships between ITGB1 and genes involved in chemoresistance and immunoresistance.

### Correlation with Related Genes

Data from GEPIA2 and GeneCardswere used to identify genes related to ITGB1. A heatmap was generated using the pheatmap package in R to visualize the correlations between ITGB1 and these related genes, providing insight into potential gene networks or pathways associated with ITGB1 in cancer. Additionally, the OS heatmap and DFS heatmap were created from GEPIA2.

## Statistical Analysis

To compare the expression differences of ITGB1 and its related genes between tumor and normal tissues across different cancer types, t-tests were used for two-group comparisons between tumor and normal tissues. For the Stage plot of ITGB1 in different cancer types, ANOVA was applied for multi-group comparisons across different cancer stages. For the survival analysis of ITGB1 and its related genes, Kaplan-Meier survival curves were generated to assess the impact of ITGB1 expression on OS and DFS, with statistical significance determined by the Log-rank test. The correlation between ITGB1 expression and chemoresistance or immunoresistance-related genes was evaluated using PCC. In survival analysis and correlation assessments, Cox proportional hazards regression models were used to adjust for potential confounding factors. The significance of the statistical findings is indicated in the figures by the following methods: one asterisk (*) or highlighted outlines for p < 0.05, and two asterisks (**) for p < 0.01.

## Conclusion

In conclusion, through bioinformatics analysis, our study highlights the significant role of ITGB1 in cancer progression, prognosis, and therapeutic resistance through comprehensive bioinformatic analyses. ITGB1 was significantly upregulated across 12 cancer types, with elevated expression correlating with advanced tumor stages and poorer survival outcomes in various cancer types. We found ITGB1 exhibited strong positive correlations with chemoresistance genes (e.g., ABCB1, CASP3) and immunoresistance markers (e.g., PD-L1, TGFB1), suggesting its dual role in mediating therapeutic evasion. Furthermore, ITGB1-associated genes (e.g., LAMC1, COL4A1) were linked to aggressive tumor behavior and poor prognosis, highlighting their collective involvement in ECM remodeling and immune modulation. These findings position ITGB1 as a promising prognostic biomarker and therapeutic target. Targeting ITGB1 or its interacting pathways could enhance the efficacy of chemotherapy and immunoresistance, particularly in tumors with high ITGB1 expression. Future studies should validate these mechanisms experimentally and explore ITGB1-targeted strategies to overcome therapeutic resistance and improve patient outcomes.

---

**Supplementary Table:**
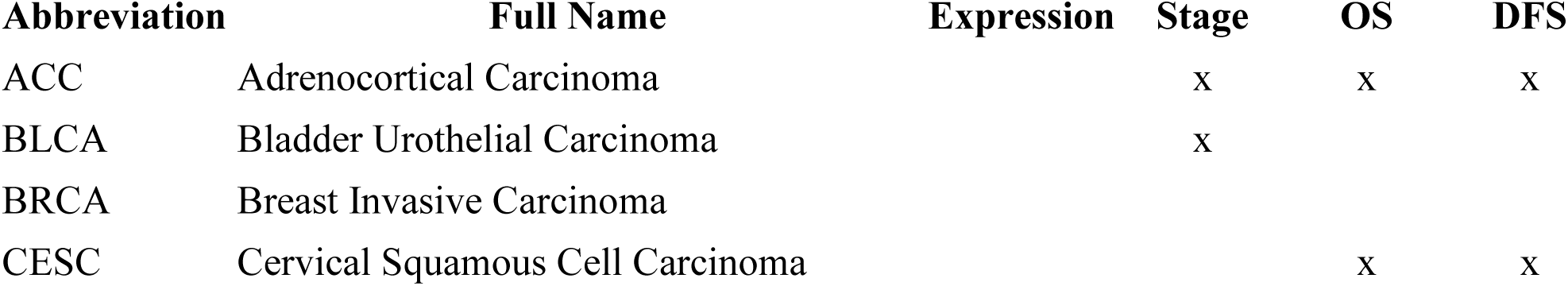
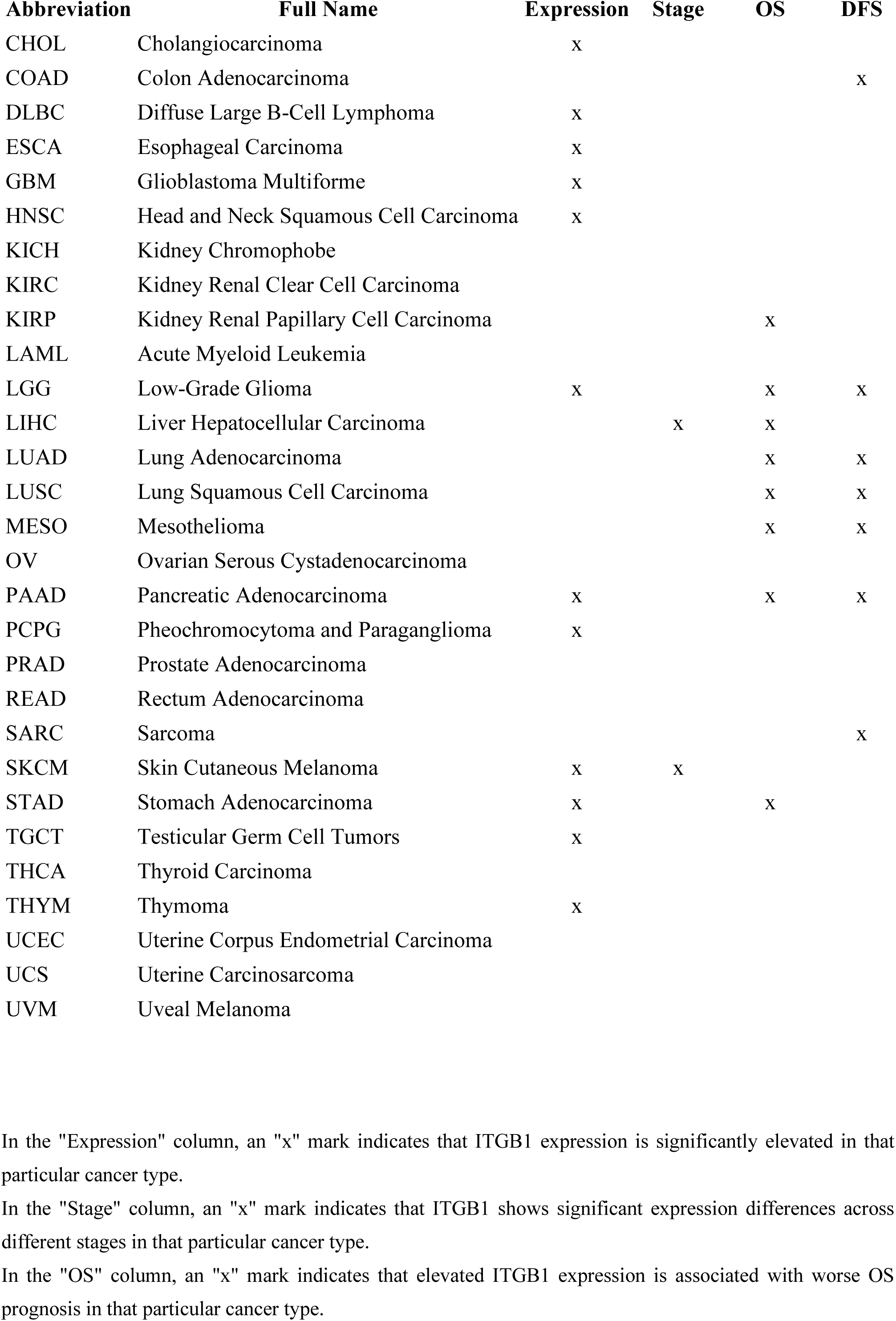
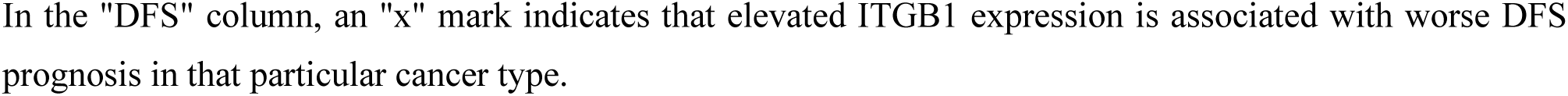
33 Cancer Types and the Summarization of ITGB1 Results

